# In silico structural analysis of EthA substitutions for ranking priority mutations leading to ethionamide resistance in *Mycobacterium tuberculosis*

**DOI:** 10.64898/2026.04.16.718980

**Authors:** Rodrigo Fernandes-Machado, Sophia Lincoln-Cardoso, Isabela Cordeiro Figueiredo, Jesus Pais Ramos, L. Caetano M. Antunes, Priscilla Goliat Capriles, Teca Calcagno Galvao

**Affiliations:** Fiocruz, Laboratório de Bacteriologia Aplicada a Saúde Única e Resistência Antimicrobiana (LabSUR), Instituto Oswaldo Cruz. Avenida Brasil 4036, Rio de Janeiro RJ 21040-361, Brazil; Fiocruz, Laboratório Nacional de Referência da Tuberculose, Centro Nacional de Referência Prof. Hélio Fraga, ENSP. Estrada da Curicica 2000, Rio de Janeiro RJ 22780-191, Brazil; Department of Computer Science, Federal University of Juiz de Fora, Juiz de Fora MG 36036-900, Brazil

**Author notes:** These authors contributed equally to the work (Rodrigo Fernandes-Machado, Sophia Lincoln-Cardoso). Laboratório de Genética Humana (LGH), Departamento de Genética e Biologia Evolutiva, Instituto de Biociências, Universidade de São Paulo. São Paulo SP 05508-090, Brazil. Department of Molecular Biosciences, Kansas University, Lawrence, USA.

**Keywords:** Ethionamide resistance, *Mycobacterium tuberculosis*, EthA, drug resistance determinant mutation

## Abstract

**Background:** Tuberculosis (TB) is the second-leading cause of deaths from infectious agents and remains a global health threat. Ethionamide (ETH) is a prodrug used in regimens for multidrug-resistant TB, and, partly due to side effects that can lead to low treatment adhesion, resistance arises. Changes in EthA, the monooxygenase that activates ETH, are the main mechanism of resistance. Yet, of hundreds of EthA substitutions found in resistant isolates, only a handful have been annotated as resistance determinants.

**Results:** An *in silico* analysis was carried out on a previously described panel of *Mycobacterium tuberculosis* clinical isolates for which genomes and ETH susceptibility testing results were available. EthA substitutions were mapped, revealing the existence of hotspots in its sequence. Visualization of the hotspots in the EthA structural model shows that they cluster in three regions, including ligand binding pockets. Models were built of twenty-three variants found in resistant isolates and changes in local configuration was mapped to identify investigate impact on ETH activation. Information from these models contributed to establishing five criteria for scoring whether substitutions are most likely to lead to resistance. Using these criteria, EthA D58G was selected and its expression is shown to increase growth in high ETH concentrations.

**Conclusion:** Functionally relevant regions of EthA are revealed and point out priority substitutions for functional studies, enhancing identification and detection of substitutions not been previously associated with resistance.

## Background

Tuberculosis (TB) is the world leading cause of deaths from a single infectious agent and the tenth leading cause of death worldwide (World Health Organization, 2024; 2025). The main cause of TB deaths is related to drug resistance, considered a public health emergency, where only 2 in every 5 reported cases receive treatment (WHO, 2024). Between 3% and 24% of cases without previous treatment history were caused by multidrug-resistant or rifampicin resistant *M. tuberculosis*, revealing extensive transmission of resistant strains (WHO, 2023). Failures in detection, access to diagnostics and treatment, and patient adherence to treatment all contribute to the emergence of resistant strains. Therefore, rapid and accurate detection of resistant *M. tuberculosis* is essential for the treatment and control of TB.

Ethionamide (ETH) is a second-line pro-drug used in the treatment of infection by multidrug-resistant strains, and can be decisive for treatment success of patients with a history of resistance (Somner & Brace, 1962; Deshpande et al., 2018). To exert its mycobactericidal effect, ETH requires an enzymatic activation step performed by EthA (Rv3854c), a NADPH- and O2-dependent Baeyer-Villiger monooxygenase (BVMO) that uses FAD as a cofactor (Vannelli et al. 2002). EthA is responsible for converting ETH into S-oxide of ethionamide, its active form. The compound inhibits the enoyl-acyl carrier protein (ACP) reductase InhA, involved in the biosynthesis of mycolic acid, an essential component of the *M. tuberculosis* cell wall (Baulard et al., 2000; DeBarber et al., 2000; Vannelli et al., 2002; Prasad et al., 2021).

Indels, premature stop codons and nonsynonymous mutations in *ethA* are the most common mechanism of resistance to ETH, followed by mutations in the promoter region of *inhA* (Vilchèze & Jacobs 2014; Banerjee et al. 1994; Morlock et al., 2003). Though other BVMOs have been shown to activate ETH (Vilchèze et al., 2008; Grant et al., 2016; Blondiaux et al., 2017), thus far evidence for their relevance in clinical resistance is limited. It has been difficult to identify specific resistance determinants among EthA substitutions, as these are often also observed in ETH susceptible (ETH-S) isolates. A dramatic example of this is the M1T/R substitution, which corresponds to a loss of function as the M1 codon is required for protein synthesis. In these strains a lack of ETH activation should result in high Minimum Inhibitory Concentration (MIC). Yet, these isolates frequently test susceptible (eg. Walker et al., 2022). Also, substitutions are found across the EthA sequence and are often found in one or few isolates. To the best of our knowledge, a single functional study has reported a causal relation between a substitution in EthA and resistance: expression of EthA-V202L (from an *M. tuberculosis* ETH resistant clinical isolate) in *Mycobacterium smegmatis* led to increased growth in presence of ETH (Anand et al., 2022).

Despite all the data linking mutations in *ethA* with resistance to ETH in *M. tuberculosis* clinical isolates, studies about the impact of substitution on EthA are limited. In this work, we evaluate the sequence and structural context of EthA substitutions from ETH resistant (ETH-R) isolates in the 2021 dataset of the WHO Catalogue of Drug Resistance Mutations (WHO 2021), to expand this previous analyses in search of resistance mutations.

## Methods

### EthA models and substitution visualization

EthA substitutions were obtained from previous reports (WHO, 2021; Walker et al., 2022). Three-dimensional models for EthA variants were generated based on a previous EthA structure (de Souza et al., 2020). The Pymol mutagenesis tool version 2.5.7 (Schrödinger, 2020) was used to substitute residues in the EthA model. The Backbone Independent Rotamers Library and the rotamer with the lowest strain value were chosen in each case. The models and their ligands were then submitted to a single round of energy minimization (500 steps in OpenBabel version 2.4.0; O’Boyle et al., 2011), the MMFF94s forcefield (Halgren, 1999), and the steepest descent algorithm. For EthA G11V, S57Y, and C137Y, a different approach was taken with two rounds of 2.500 steps each using the General Amber Forcefield (Wang et al., 2004). Finally, 2D interaction maps were created for the resulting models using Maestro version 13.6 (Schrödinger, 2023) and PyMOL version 2.5.7 (Schrödinger, 2020). Electrostatic potential maps were calculated using the Adaptive Poisson-Boltzmann Solver (APBS) PyMOL plugin (Jurrus et al., 2018; Schrodinger, 2020), with charges ranging between –10^4^ and 10^4^ eV. pKa values were predicted using the PropKa on-line server version 2.0 (Li et al., 2005).

### Bacterial cells and cloning

For construction of pMAR, Gibson cloning was used to introduce the *Mycobacterium bovis* BCG *ethA_*P*ethAR_ethR* locus into the KpnI site of pUS972 (Norazmi et al., 1999). Mutations in pMAR for expressing variants M1T, S57Y, and D58G were obtained by site directed mutagenesis (SDM) following the SPRINP protocol (Edelheit et al., 2009). Overlapping oligonucleotides were synthesized with a length of 31-35 nucleotides, with the desired mutation in the center.

Independent amplification reactions were prepared in two tubes, each containing: pMAR (∼250 ng), primer (forward or reverse) at 0.5 μM, Phusion™ GC Buffer 1x (Thermo), 0.2 mM dNTPs, Phusion™ High-Fidelity DNA polymerase 0.02 U/μL (Thermo), and 3% (v/v) DMSO in a final volume of 25 μL. The reactions were combined, subjected to denaturation (95°C for 5 minutes) and sequential cooling (90°C and 80°C for 1 minute, 70°C, 60°C, 50°C and 40°C for 30 seconds). Following digestion with DpnI (10 U/μL) and inactivation at 80°C for 20 minutes, the reactions were dialysed against water in nitrocellulose filters (pore size 0.025 um; Millipore). The reactions were then transformed into *Escherichia coli* One Shot TOP10 (Invitrogen). The *ethA-ethR* locus from kanamycin resistant colonies was amplified by colony PCR and clones with the mutations were identified by Sanger sequencing.

### Susceptibility testing for ethionamide in *M. smegmatis*

Competent *Mycobacterium smegmatis* cells were transformed by electroporation with 0.5 ug of plasmids. Cells were plated on 7H10 medium supplemented with 10% (v/v) ADC, containing 25 μg/mL of kanamycin, and incubated for 48 hours at 37°C. ETH (Sigma) was dissolved at a concentration of 16 mg/mL in ethylene glycol (Sigma) with heating ∼40°C and vortexing until completely dissolved. The stock was diluted in Milli-Q water to 8 mg/mL and filtered ( MF-Millipore™ MCE 0.22 μm filter). Colonies on the transformation plates were collected with a loop and placed in a glass tube containing glass beads and 500 μL of water. The mixture was vortexed (30 to 60 seconds), followed by addition of 5 mL of water and a 5 minute incubation. Drops from the region below the meniscus of the suspension were transferred to a new tube containing 2 mL of saline until the turbidity reached 0.5 on the McFarland scale, (with Vitek DensiChek bioMérieux). Serial dilutions from the standard to 10^−6^ were prepared by adding 10 ul of sample to 90 ul of 7H9 supplemented with ADC, 0.05% (v/v) TWEEN 80, and 25 μg/mL of kanamycin.

Drops of 5 µL were seeded in fresh plates containing 30 mL of 7H10 medium supplemented with ADC, kanamycin at 25 μg/mL and ethionamide at 0–80 μg/mL, followed by incubation at 37°C for 72 h.

## Results

### Substitutions in EthA represent a significant proportion of changes found in ETH-R isolates

In 2021, the publication of the WHO-endorsed catalog of *M. tuberculosis* mutations associated with resistance provided a global standard. By compiling whole genome sequencing and drug susceptibility testing data of a large and diverse set of isolates (∼38,000), the study generated a catalog of mutations with data for 13 anti-TB drugs, including ETH (WHO, 2021; Walker et al., 2022).

Mutations were designated as solo R or solo S when, in a given isolate, no other changes were identified in the genes analyzed for a specific drug (Walker et al., 2022). Variants were considered as resistance determinants based on criteria such as frequency, odds ratios (ORs), positive predictive value (PPV), and being within the 95% confidence interval (CI) (Walker et al. 2022). In the ETH data, 1438 changes in *ethA-ethR*, *fabG1-inhA* and promoter, *mshA*, *ndh* and *rv3083* were identified in 13,620 isolates. Of these, 365 occur in solo R isolates. The most frequently found types of changes in ETH solo R isolates were in the *inhA* promoter (739 isolates), *ethA* nonsynonymous substitutions (491 isolates) or indels and premature stop codons (425 isolates). That *ethA* nonsynonymous mutations are found in 29% of ETH-R isolates bearing solo R mutation highlights their significant contribution to ETH resistance. Yet, only ten EthA substitutions were ranked as associated with resistance or as interim associated (M1T, M1R, G11V, Y32D, S57Y, T88I, A341V, R207G, P378L, S390F; Walker et al. 2022).

### Distribution of substitutions across EthA primary sequence and identification of hotspots

Given the major role of EthA in resistance, it is important to investigate which are likely to impair ETH activation by impacting residues relevant in structure and/or function. Thus, to further explore this large data set, we first mapped solo R mutations/substitutions from ETH-R and ETH-S isolates in the EthA primary sequence (**Figure 1**). In the 491 ETH-R isolates bearing EthA substitutions as solo R mutation, 171 substitutions were distributed across 126 amino acid positions (**Table S1**; **Figure S1)** and a subset of neighboring residues concentrated a high number of solo R mutations. When at least 5 ETH-R isolates with solo R EthA substitutions were mapped to these short stretches of amino acids, we referred to them as hotspots. The hotspots are also visible by plotting the solo R EthA substitutions data from Walker and colleagues (2022) (**Figures S1 and S2**).

**Figure 1.**
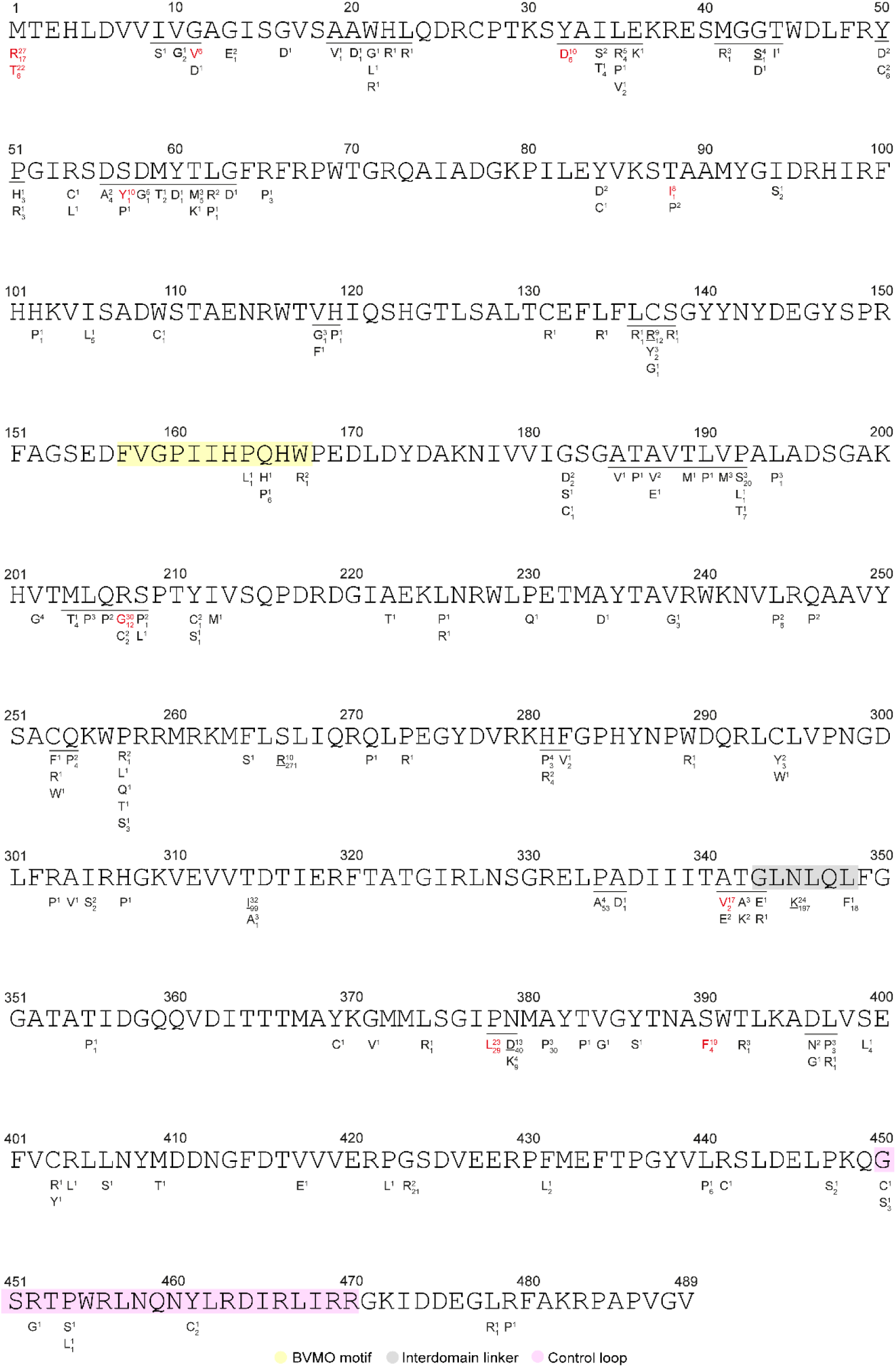
Distribution of solo mutations in EthA. The number of isolates with a given substitution in ETH-R and ETH-S isolates are shown as super- and subscript, respectively. Regions whose substitutions were found in at least five resistant isolates (hotspots) are underlined. Substitutions associated with resistance by the WHO analysis are shown in red. EthA regions are shown as colored boxes.

### Hotspots of EthA substitutions form spatially distributed clusters

Next, we mapped the hotspot residues to the EthA model in complex with NAD, FAD and ETH (de Souza et al., 2020). Hotspots are in spatial proximity across three regions, which we refer to as clusters 1, 2, and 3 (**Figure 2A**). Hotspots from across the sequence compose clusters 1 and 2 (including four resistance determinant variants: G11D, Y32D, R207G, A341V) and cluster 3 comprises a single stretch of amino acids (D56-G63) containing the resistance determinant S57Y (**Figure 2B**).

**Figure 2:**
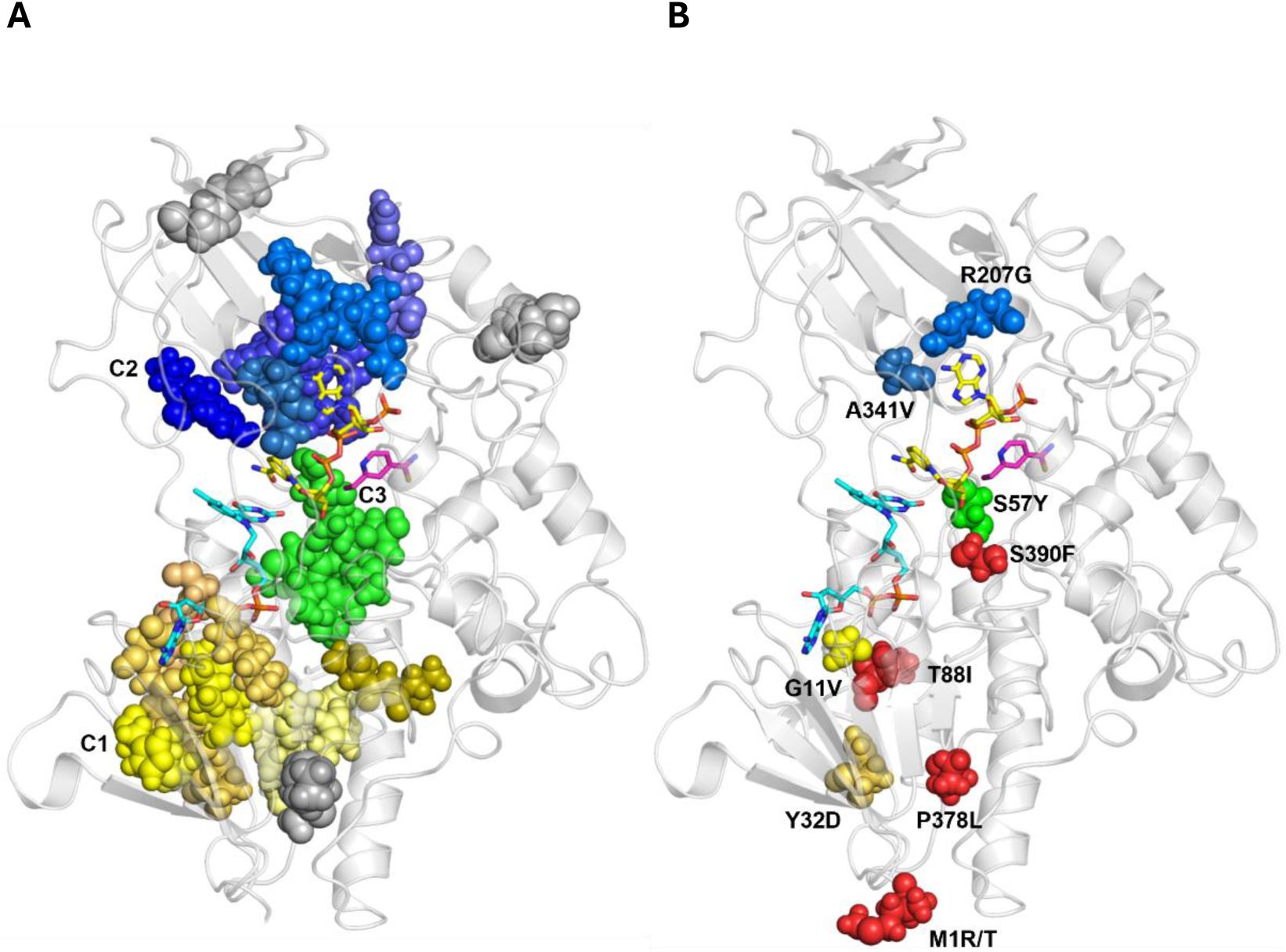
Residues in resistance hotspots form clusters in the 3D EthA model. (A) Residues in hotspots are shown as spheres. Hotspots that cluster are colored in shades of yellow (cluster C1), blue (cluster C2) and green (cluster C3). Hotspots that do not cluster are shown in gray. (B) Residues whose substitution is associated with resistance in the WHO Catalogue are highlighted in colors corresponding to their respective clusters. Those that are off clusters are shown as red spheres.

Notably, clusters 1 and 2 are made up of hotspots that are not contiguous in the primary sequence (**Figure 3**; **Table 1**). Substitutions M1R/T, T88I, P378L and S390F were catalogued as resistance determinants, but are found off clusters.

**Figure 3.**
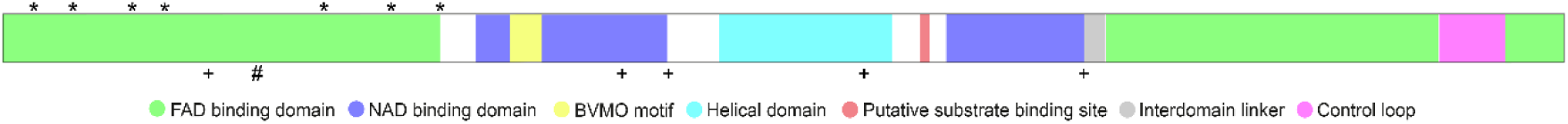
Hotspots contributing to clusters are spread over the EthA sequence. Schematic representation of EthA’s primary structure displaying putative domains and hotspots positions. Hotspots in clusters 1, 2 and 3 are marked with ★, **+**, and **#,** respectively.

**Table 1.**
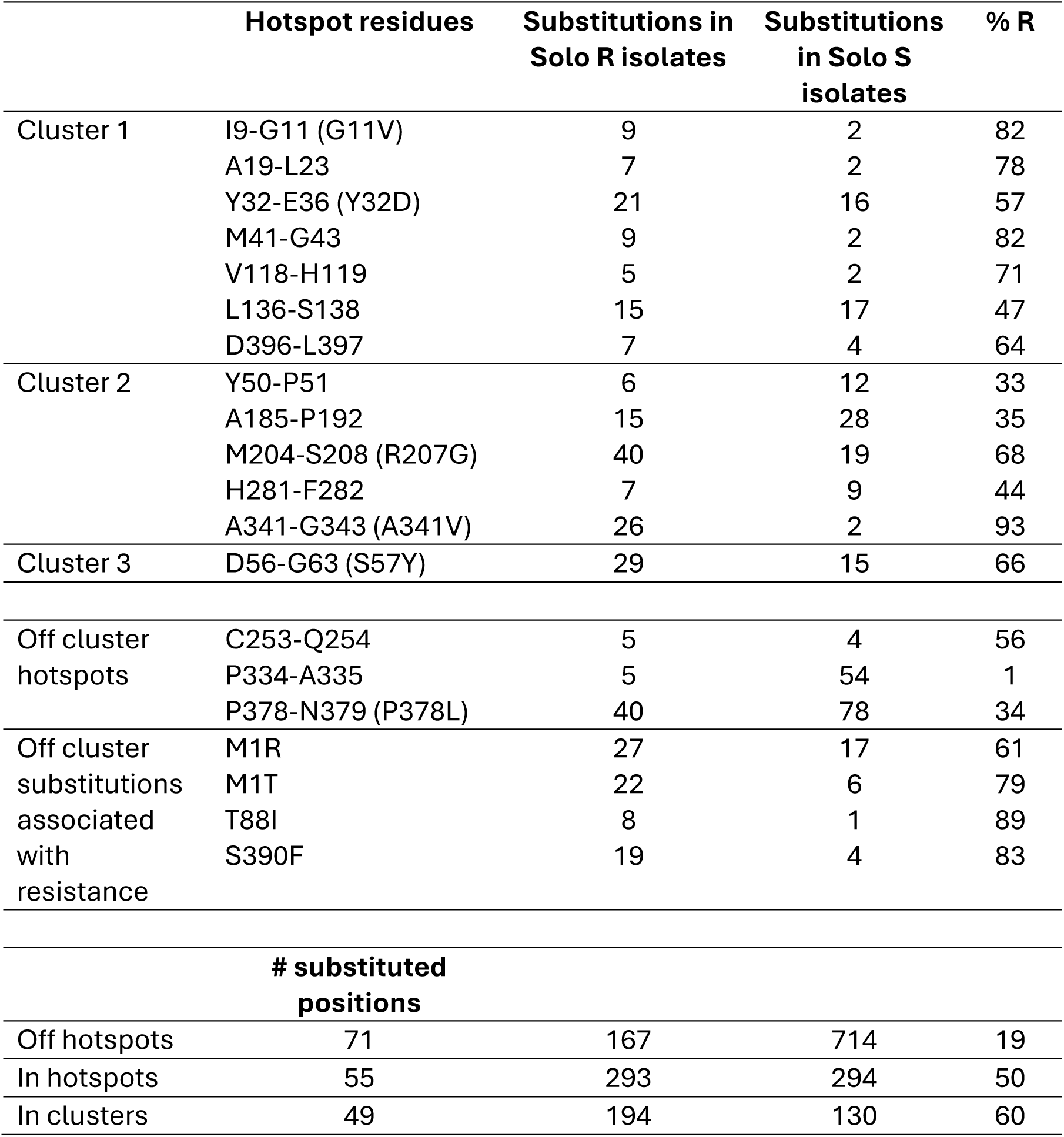
Substitutions hotspots in EthA concentrate solo R *ethA* mutations of ETH-R isolates. Substitutions associated with resistance found in clusters are shown in parenthesis.

The significance of the hotspots is made evident by the observation that only 19% of isolates bearing EthA substitution off hotspots are resistant (**Table 1**), whereas this proportion rises to 50 and 60% for isolates bearing substitutions in hotspots and clusters, respectively. Also, in nine of thirteen hotspots, most isolates bearing substitutions are resistant. For example, of 28 isolates with solo R mutations in hotspot A341-G343, 26 are ETH-R. Thus, hotspots concentrate substitutions in resistant isolates, suggesting that they are a map to resistance determinants, and that substitutions outside them are more likely to represent sequence diversity not associated with ETH selective pressure. This finding reduces from 126 to 55 the number of positions to be considered priority when looking for EthA variants with a functional role in resistance.

### Evaluating the impact of selected EthA substitutions

In the ETH dataset (Walker etal., 2021), there were 171 EthA substitutions found as solo R mutation in ETH-R isolates, spread over 126 positions (**Figure 1**). Of these, only thirty-five were identified in at least 3 solo R isolates (**Table 2**). A few substitutions were identified in over fifty isolates (S266L, H281P, T314I, P334A, N345K, P378L, N379D, A381P), the majority of which tested ETH-S, suggesting that these mutations are frequent alleles natural do *ethA* genetic diversity that do not modulate resistance.

**Table 2.** Substitutions found in three or more ETH-R isolates as solo R mutations. Data adapted from Walker et al., 2022.

| Substitution | Cluster | SOL<br>O<br>R | SOLO<br>S | #<br>isolates | % solo<br>R/solo RS | % solo<br>R/R | Non solo<br>R |
| --- | --- | --- | --- | --- | --- | --- | --- |
| T314I<sup>#</sup> | - | 32 | 99 | 131 | 23 | 80 | 8 |
| R207G<sup>*</sup> | 2 | 30 | 12 | 42 | 71 | 100 | 0 |
| M1R<sup>*</sup> | - | 27 | 17 | 44 | 61 | 100 | 0 |
| N345K | - | 24 | 197 | 317 | 8 | 22 | 87 |
| P378L<sup>*</sup> | - | 23 | 29 | 56 | 41 | 92 | 2 |
| M1T<sup>*</sup> | - | 22 | 6 | 30 | 73 | 92 | 2 |
| S390F<sup>*</sup> | - | 19 | 4 | 23 | 83 | 100 | 0 |
| A341V<sup>*</sup> | 2 | 17 | 2 | 20 | 85 | 94 | 1 |
| N379D<sup>#</sup> | - | 13 | 40 | 53 | 25 | 100 | 0 |
| Y32D<sup>*</sup> | 1 | 10 | 6 | 16 | 63 | 100 | 0 |
| S57Y<sup>*</sup> | 3 | 10 | 1 | 12 | 83 | 91 | 1 |
| S266R | - | 10 | 271 | 474 | 2 | 6 | 145 |
| C137R<sup>#</sup> | 1 | 9 | 12 | 23 | 39 | 82 | 2 |
| T88I<sup>*</sup> | - | 8 | 1 | 9 | 89 | 100 | 0 |
| G11V<sup>*</sup> | 1 | 6 | 0 | 7 | 86 | 86 | 1 |
| D58G<sup>#</sup> | 3 | 5 | 1 | 6 | 83 | 100 | 0 |
| L35R<sup>#</sup> | 1 | 5 | 4 | 11 | 45 | 71 | 2 |
| G43S<sup>#</sup> | 1 | 4 | 1 | 5 | 80 | 100 | 0 |
| V202G<sup>#</sup> | - | 4 | 0 | 4 | 100 | 100 | 0 |
| H281P | 2 | 4 | 3 | 160 | 3 | 3 | 131 |
| P334A | - | 4 | 53 | 66 | 6 | 36 | 7 |
| N379K | - | 4 | 7 | 13 | 31 | 100 | 0 |
| M41R<sup>#</sup> | 1 | 3 | 1 | 4 | 75 | 100 | 0 |
| T61M<sup>#</sup> | 3 | 3 | 5 | 8 | 38 | 100 | 0 |
| V118G<sup>#</sup> | 1 | 3 | 1 | 4 | 75 | 100 | 0 |
| C137Y<sup>#</sup> | 1 | 3 | 2 | 6 | 50 | 75 | 1 |
| V191M<sup>#</sup> | 2 | 3 | 0 | 3 | 100 | 100 | 0 |
| P192S | 2 | 3 | 20 | 25 | 12 | 60 | 2 |
| L194P | - | 3 | 1 | 4 | 75 | 100 | 0 |
| L205P | 2 | 3 | 0 | 3 | 100 | 100 | 0 |
| T314A<sup>#</sup> | - | 3 | 1 | 7 | 43 | 50 | 3 |
| T342A | 2 | 3 | 0 | 4 | 75 | 75 | 1 |
| A381P | - | 3 | 30 | 196 | 2 | 4 | 70 |
| T392R | - | 3 | 1 | 4 | 75 | 100 | 0 |
| L397P | 1 | 3 | 3 | 6 | 50 | 100 | 0 |

\*Annotated as resistance determinants (Walker et al., 2022). <sup>#</sup>substitution chosen for further evaluation. % solo R/R: percentage of solo R isolates out of all R isolates. Non solo R: number of R isolates bearing additional variants in *ethA* or other ETH genes.

To further evaluate the impact of EthA substitutions, twenty-three were chosen for further evaluation: ten substitutions found mostly in ETH-R isolates, eight resistance determinant substitutions (Walker et al., 2022; WHO, 2021), and five substitutions found mostly in ETH-S isolates. Models for each protein were built based on EthA_14053 structural model (de Souza et al., 2020), with the protein mutagenesis wizard tool (PyMol v2.5.5). The EthA_14053 model was chosen as it had the EthA ligands docked, and in this conformation, obtained through MD simulations, the control loop and the interdomain linker are proximal, as described to occur during BVMO catalysis (Mirza et al., 2009).

To evaluate the potential impact of these twenty-three substitutions on protein-ligand interactions, each model was optimized and the distances to the three ligands (NAD, FAD and ETH) were measured in Pymol. Eight positions are at ≤4 Å of NAD or FAD (L35R, G43S, C137Y/R and resistance determinants G11V, S57Y, R207G and S390F), showing a reliable distance for a direct molecular interaction (**Figure 4**). Notably, all of these except S390F are in clusters. Substituted amino acids were not in proximity to ETH, with the closest being at 8.4Å (S390F). Note that of the twenty-three substitutions chosen for analysis, fifteen are in clusters. Thus, that seven of these are in close vicinity of FAD and NAD points to two important aspects. First, these data further support the quality of the EthA model. Second, the colocalization of a high proportion of amino acids within close range of ligands strengthen the concept and importance of the clusters. This is made evident as positions A341V and M41R, within 6 Å of NAD and FAD, respectively, are also in clusters. Such proximity suggests that reduced ETH activation is a consequence of impaired EthA-ligand interactions and-or catalysis.

**Figure 4:**
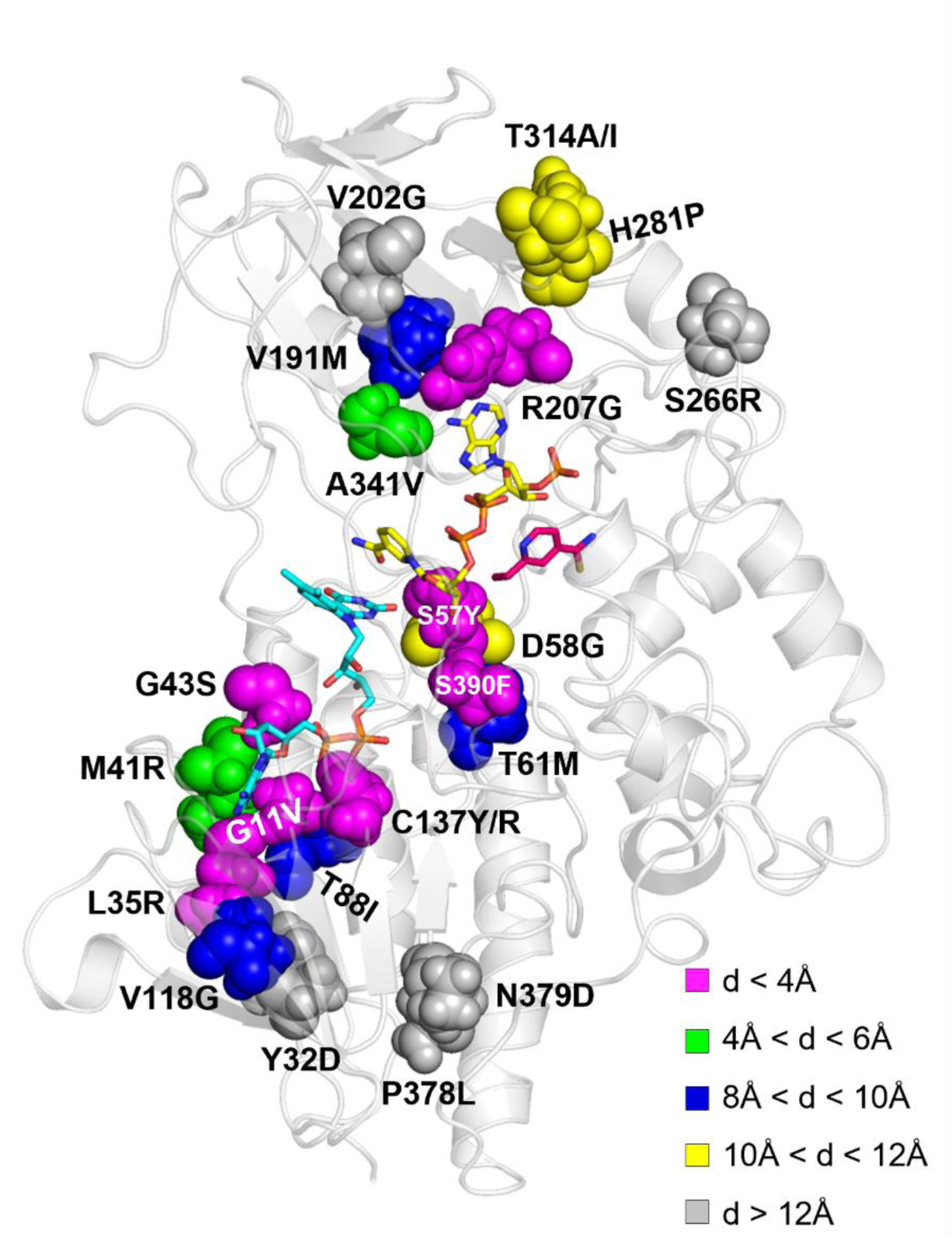
Proximity of EthA ligands to the twenty-three amino acid substitutions selected for analysis. Residues (spheres) are colored according to their distance (d) to the nearest ligand (sticks). FAD is shown in light blue, NAD in yellow and ETH in pink.

### Impact of substitutions in EthA and ligand interactions

The twenty-three optimized structures were used in Pymol and Maestro to map interactions and generate 2D maps of FAD and NAD, and PropKa was used to evaluate changes in pKa and electrostatic potential (**Figure5; Figures S3, S4 and S5**). The results are summarized in **Table 3**, showing the characteristics detected for each substitution. Changes in the FAD and NAD interaction network were detected in five structures: G11V, G43S, S57Y, C137Y/R, and R207G.

**Table 3:**
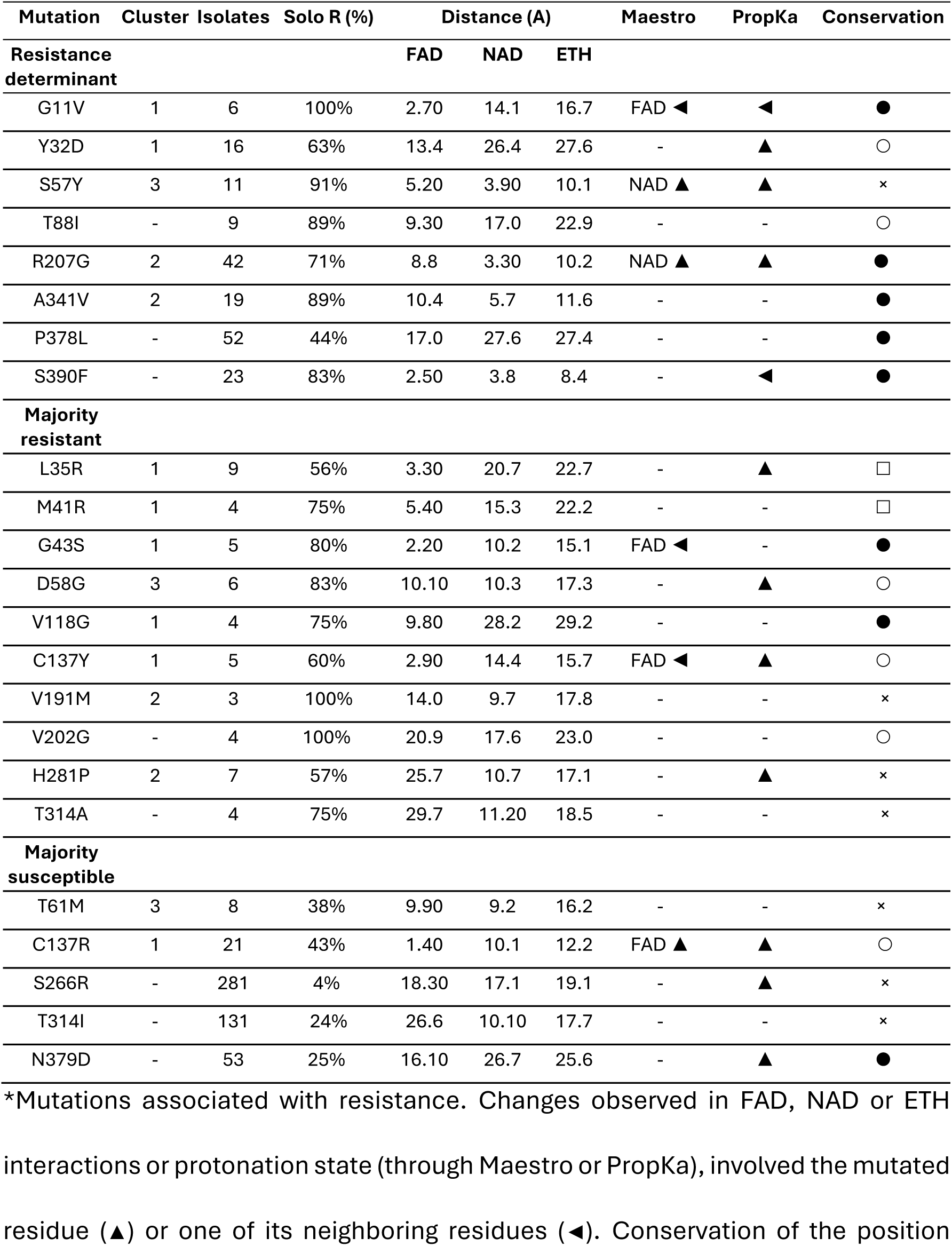
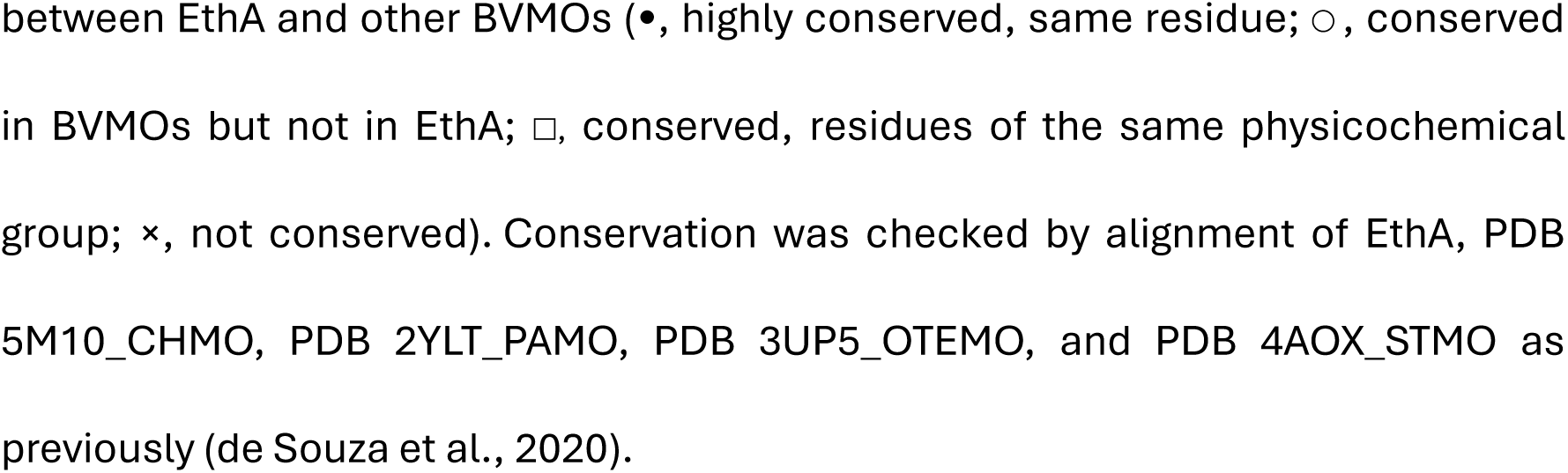
Summary of the features in the EthA substitutions chosen for analysis. The cluster, number of solo R isolates, percentage of solo R (over solo RS) isolates, distance from ligands, Maestro and PropKa analysis, and sequence conservation are shown for each of the twenty-three selected substitutions.

| Mutation | Cluster | Isolates | Solo R (%) | Distance (Å) |  |  | Maestro | PropKa | Conservation |
| --- | --- | --- | --- | --- | --- | --- | --- | --- | --- |
| Resistance determinant |  |  |  | FAD | NAD | ETH |  |  |  |
| G11V | 1 | 6 | 100% | 2.70 | 14.1 | 16.7 | FAD ◀ | ◀ | ● |
| Y32D | 1 | 16 | 63% | 13.4 | 26.4 | 27.6 | - | ▲ | ○ |
| S57Y | 3 | 11 | 91% | 5.20 | 3.90 | 10.1 | NAD ▲ | ▲ | × |
| T88I | - | 9 | 89% | 9.30 | 17.0 | 22.9 | - | - | ○ |
| R207G | 2 | 42 | 71% | 8.8 | 3.30 | 10.2 | NAD ▲ | ▲ | ● |
| A341V | 2 | 19 | 89% | 10.4 | 5.7 | 11.6 | - | - | ● |
| P378L | - | 52 | 44% | 17.0 | 27.6 | 27.4 | - | - | ● |
| S390F | - | 23 | 83% | 2.50 | 3.8 | 8.4 | - | ◀ | ● |
| Majority resistant |  |  |  |  |  |  |  |  |  |
| L35R | 1 | 9 | 56% | 3.30 | 20.7 | 22.7 | - | ▲ | □ |
| M41R | 1 | 4 | 75% | 5.40 | 15.3 | 22.2 | - | - | □ |
| G43S | 1 | 5 | 80% | 2.20 | 10.2 | 15.1 | FAD ◀ | - | ● |
| D58G | 3 | 6 | 83% | 10.10 | 10.3 | 17.3 | - | ▲ | ○ |
| V118G | 1 | 4 | 75% | 9.80 | 28.2 | 29.2 | - | - | ● |
| C137Y | 1 | 5 | 60% | 2.90 | 14.4 | 15.7 | FAD ◀ | ▲ | ○ |
| V191M | 2 | 3 | 100% | 14.0 | 9.7 | 17.8 | - | - | × |
| V202G | - | 4 | 100% | 20.9 | 17.6 | 23.0 | - | - | ○ |
| H281P | 2 | 7 | 57% | 25.7 | 10.7 | 17.1 | - | ▲ | × |
| T314A | - | 4 | 75% | 29.7 | 11.20 | 18.5 | - | - | × |
| Majority susceptible |  |  |  |  |  |  |  |  |  |
| T61M | 3 | 8 | 38% | 9.90 | 9.2 | 16.2 | - | - | × |
| C137R | 1 | 21 | 43% | 1.40 | 10.1 | 12.2 | FAD ▲ | ▲ | ○ |
| S266R | - | 281 | 4% | 18.30 | 17.1 | 19.1 | - | ▲ | × |
| T314I | - | 131 | 24% | 26.6 | 10.10 | 17.7 | - | - | × |
| N379D | - | 53 | 25% | 16.10 | 26.7 | 25.6 | - | ▲ | ● |

\*Mutations associated with resistance. Changes observed in FAD, NAD or ETH interactions or protonation state (through Maestro or PropKa), involved the mutated residue (▲) or one of its neighboring residues (◀). Conservation of the position
between EthA and other BVMOs (●, highly conserved, same residue; ○, conserved in BVMOs but not in EthA; □, conserved, residues of the same physicochemical group; ×, not conserved). Conservation was checked by alignment of EthA, PDB 5M10\_CHMO, PDB 2YLT\_PAMO, PDB 3UP5\_OTEMO, and PDB 4AOX\_STMO as previously (de Souza et al., 2020).

The positions substituted in G11V, G43S and C137R/Y are near each other in the EthA model, lining the pocket of the adenine nucleotide portion of FAD. The solo R mutation G11V was found exclusively in resistant isolates (six in total) and was associated with resistance by Walker and colleagues (2022). G11 is highly conserved, and its substitution by a bulky and hydrophobic valine is concomitant with a small change in the position of FAD, as well as placing V11 within 3 Å of FAD. G11 is just 2.7Å from FAD, therefore being part of its interaction region, in addition to being conserved and close to other conserved residues (A12, G13, G16). The 2D map of the G11V structure shows loss of two hydrogen bonds between FAD and amino acids G13 and C137, and one new bond, to K37 (**Figure S3**). Furthermore, the impact of the substitution was also evaluated with PropKa, showing loss of hydrogen bonding between the side chains of C137 and T383, and of interaction with the backbone between C137 and G13. Electrostatic potential maps did not show visible difference between G11 and V11 (**Figure S5**).

C137 is conserved in BVMOs but not in EthA, and is in the immediate vicinity of FAD; C137Y and C137R are 2.9 and 1.4Å from FAD, respectively. The 2D interaction map shows the bond network in this region is affected in both R137 and Y137 (**Figure S3**). Y140 stands between position 137 and FAD, imposing steric limitations. In the EthA model, C137 hydrogen bonds to a hydroxyl group in FAD’s phosphate; this bond is lost in the C137Y structure, while in C137R it is substituted by two new interactions with the other FAD phosphate group. Loss of backbone hydrogen bonds with G13, S15, and G16 was also suggested for both. An increase in pKa values was predicted through PropKa for both Y137 and R137 mutant structures (16,5% and a 19,74%, respectively), reflecting the change from the polar uncharged short cysteine to the hydrophobic tyrosine or basic arginine (**Figure 5; Figure S3**). Yet, while 60% of isolates bearing the C137Y are ETH-R, this goes down to 43% for the C137R substitution. Such examples point to individual substitutions having varying impact on MIC, suggesting a degree of flexibility in that region and examplifying the complexities involved in pin pointing resistance mechanisms.

**Figure 5.**
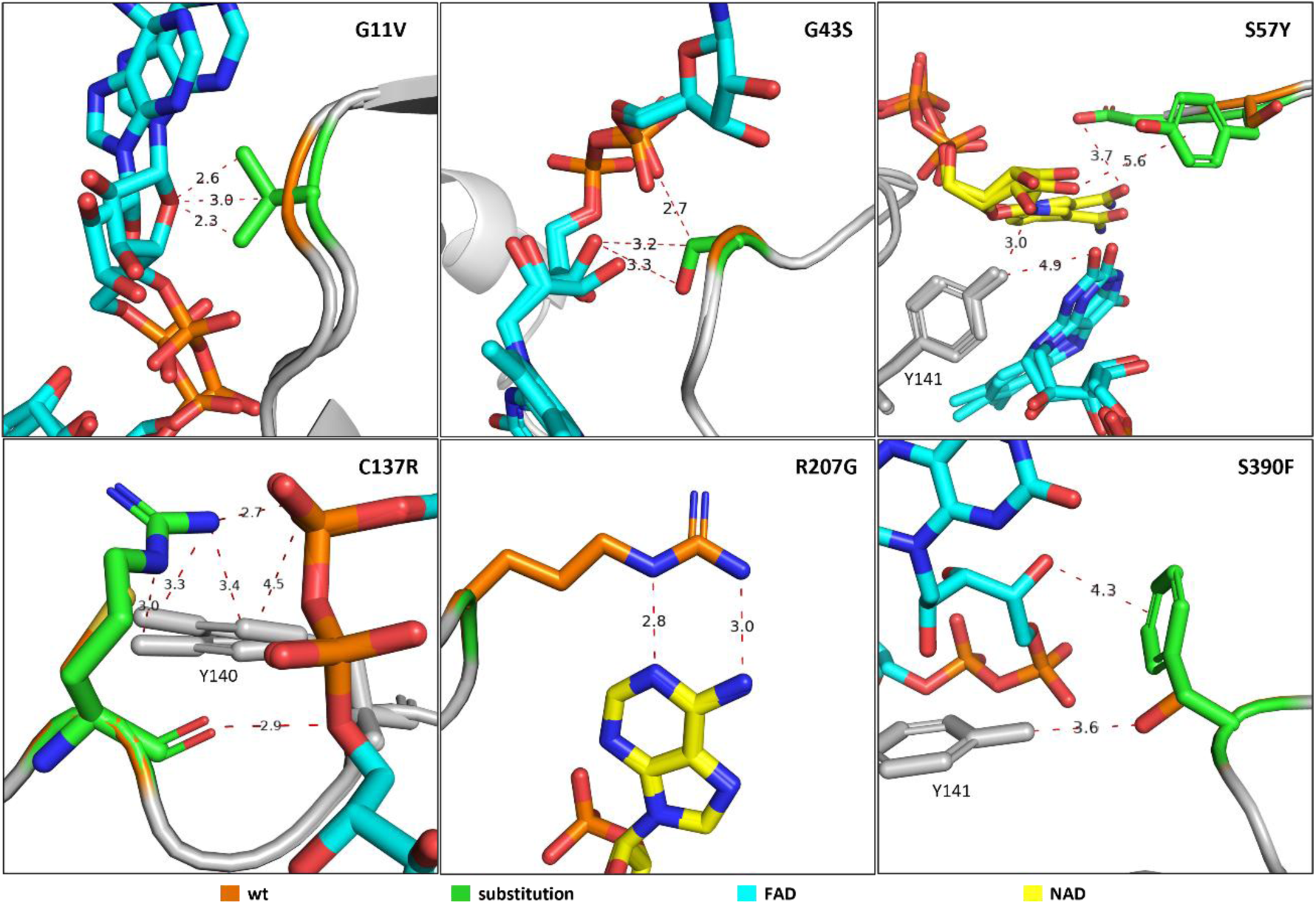
Superposition of wild-type and models in which the substitution resulted in changes in the ligand interaction pattern as judged by Maestro and/or Pymol. **I**mages and measurements (dotted lines) generated using PyMOL, through superposition of the wild-type structure (EthA 14053) and each mutant individually. Orange, wt residues; green, substitutions; blue, FAD; yellow, NAD. Hydrogens and main chain atoms not involved in interactions are omitted for better visualization.

G43 is a conserved residue in BVMOs, in EthÁs OTEMO-like region (da Silva et al., 2018). Though G43S was not annotated as a resistance determinant, of five isolates carrying this substitution, four were resistant. The change to serine places the polar side chain within 3Å of FAD, but no hydrogen bond is detected in the 2D map of the structure; rather, there is a loss of the FAD bond to the neighbouring T44 (**Figure S3**).

Substitutions S57Y and S390F are resistance determinants, and these positions are in the region where the NAD adenine nucleotide and the FAD flavin mononucleotide approach each other (**Figure 4**). S57 is not conserved in BVMOs and in the S57Y structure a new hydrogen bond is observed in the 2D map, between Y57 and the ribose of NAD. It is possible that new coulomb interactions between Y57 and Y141 and between Y57 and D56, along with the loss of a side-chain hydrogen bond between D58 and S57 affect the transfer of electrons with NAD, perhaps by imposing steric inhibition due to the aromatic rings, leading to a decrease in protein mobility. These, together with a change in pKa detected by PropKa, suggest how activation of ETH may be affected by the local perturbation to bond network and rigidity. Changes in pKa are also detected for S390F, a conserved position also near FAD (2.5A).

R207 is 3.3A from the NAD nicotinamide portion and partially buries the cofactor in the enzyme (**Figure S4**). In the R207G structure, with the removal of the large arginine side chain, this portion of NAD becomes solvent exposed. R207 is highly conserved, pointing to an important role in ETH activation.

### Ranking mutations for priority in functional characterization

Currently, ETH DST results coupled with sequencing resistance genes do not yield a positive predictive power to confidently assign EthA substitutions as resistance determinants. Due to the large variety of EthA isolates and the complexity of functional studies, a set of five criteria is proposed for directing the choice of substitutions for further characterization. These are: presence in spatial clusters in EthA, whether the majority of tested isolates are solo ETH-R, being within 6 A of ligands, conservation in homologs, and whether the substitution impacted on the bond pattern around the residue and-or with nearby ligands (**Table 4**). Of eight resistance determinants assigned by Walker and colleagues (2022), five score in 4 or 5 criteria (63%); this proportion is 50% among substitutions found as solo R in at least three isolates, with five of ten substitutions scoring in 4 or 5 criteria. No substitutions found in a majority of solo S isolates scored in 4 or 5 criteria. Thus, besides the necessary characterization of the annotated resistance determinants, the data presented here reveals that substitutions L35R, M41R, G43S, D58G, and C137Y as priority in further studies.

**Table 4.** Summary of detected characteristics of select EthA substitutions.

| Substitution | Cluster | Majority solo R | Ligand < 6A | Conservation | Changes in bond pattern | Attends to criteria |
| --- | --- | --- | --- | --- | --- | --- |
| G11V | 1 | y | y | ● | y | 5 |
| Y32D | 1 | y | n | ○ | y | 4 |
| S57Y | 3 | y | y | × | y | 4 |
| T88I | - | y | n | ○ | n | 2 |
| R207G | 2 | y | y | ● | y | 5 |
| A341V | 2 | y | n | ● | n | 3 |
| P378L | - | n | n | ● | n | 1 |
| S390F | - | y | y | ● | y | 4 |
|  | 5/8 (63%) | 7/8 (88%) | 4/8 (50%) | 7/8 (88%) | 5/8 (63%) | 5/8 (63%) |
| **Majority resistant** | **8** | **10** | **4** | **7** | **5** |  |
| L35R | 1 | y | y | □ | y | 5 |
| M41R | 1 | y | y | □ | n | 4 |
| G43S | 1 | y | y | ● | y | 5 |
| D58G | 3 | y | n | ○ | y | 4 |
| V118G | 1 | y | n | ● | n | 3 |
| C137Y | 1 | y | y | ○ | y | 5 |
| V191M | 2 | y | n | × | n | 2 |
| V202G | - | y | n | ○ | n | 2 |
| H281P | 2 | y | n | × | y | 3 |
| T314A | - | y | n | × | n | 1 |
|  | 8/10 (80%) | 10/10 (100%) | 4/10 (40%) | 7/10 (70%) | 5/10 (50%) | 5/10 (50%) |
| **Majority susceptible** |  |  |  |  |  |  |
| T61M | 3 | n | n | × | n | 1 |
| C137R | 1 | n | y | ○ | y | 3 |
| S266R | - | n | n | × | n | 0 |
| T314I | - | n | n | x | n | 0 |
| N379D | - | n | n | ● | n | 1 |
|  | 2/5 (40%) | 0/5 (0%) | 1/5 (20%) | 2/5 (40%) | 1/5 (20%) |  |

To start testing these predictions, EthA variant D58G was expressed in *M. smegmatis* and the ability to grow in increasing ETH concentrations was tested. As shown in Figure 6, EthA D58G is strongly resistant to ETH, growing in the highest concentration tested (80 ug/ml). Expression of two resistance determinants (M1T and S57Y) yielded similar results, while expression of wild type EthA or empty vector did not support growth as ETH increased. D58G satisfies four of five criteria for prioritasing variants, and this result validates their use in ranking priority substitutions.

**Figure 6.**
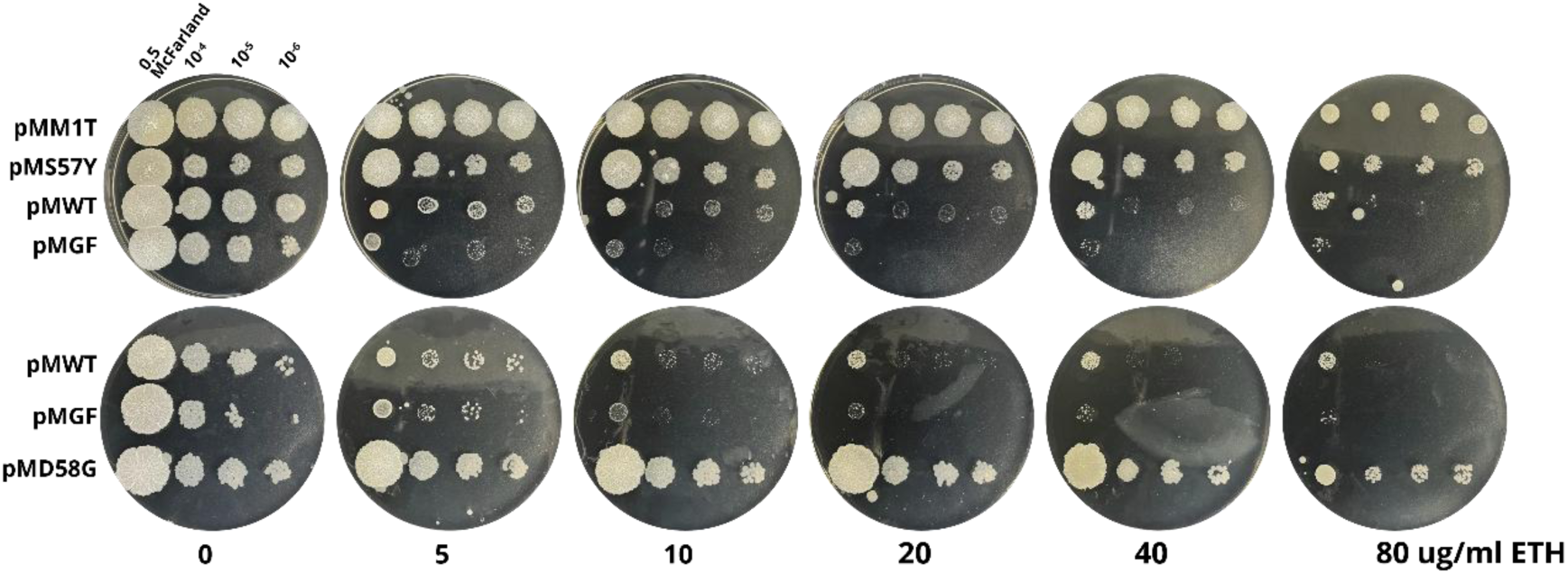
EthA D58G confers resistance to ETH in a *M. smegmatis* complementation assay. *M. smegmatis* carrying empty plasmid (pMGF) or plasmid with the *M. tuberculosis* H37Rv *ethA-ethR* locus (wild type or coding for the M1T, S57Y or D58G substitutions) was diluted to 0.5 McFarland and serial dilutions were plated in 7H10 agar with increasing concentrations of ETH.

## Discussion

The search for resistance determinants by phenotype-genotype became more sophisticated as DST with the MGIT 960 automated system became more common (reviewed in Rodrigues et al., 2008), followed by the expansion in WGS data acquisition and analysis. A hallmark came with the first catalogue of mutations, using a training dataset that allowed predictions to be made based on statistical analysis (Walker et al., 2015). The WHO catalogue of resistance mutations represents a fundamental reference, and its reach has been built on. For instance, it has been used to map regional or national isolate profiles more precisely (Pei et al., 2024). Also, modification of the WHO algorithm and inclusion of data from additional isolates yielded twenty-three substitutions annotated as “associated with resistance – interim,” implicating these as worthy of further evaluation (Kulkarni et al., 2025). Importantly, sixteen of these are within the clusters identified in this work, including D58G, shown to confer growth in high concentrations of ETH. Thus, this work can complement and validate the algorithm assignments. These results contribute to increased sensitivity of molecular diagnostic tests. An example is Deeplex Myc-TB targeted NGS for resistance detection from sputum and culture (Jouet et al., 2021), which is becoming available in reference laboratories and includes *ethA* sequencing. As ∼25% of ETH-R isolates carry *ethA* nonsynonymous mutations, expanding the number of substitutions validated as resistance determinants is necessary to improve the sensibility of Deeplex Myc-TB, which currently is very limited.

The analysis of twenty-three substitutions hints at different ways in which mutations can lead to resistance, namely proximity to FAD or NAD, changes around ligands and protonation states. For example, a comparison between EthA substitutions P378L and N379D can illustrate how the current analysis can yield insights into DST results. P378L was in the 2021 WHO Catalogue, found in 23 and 29 solo R and solo S isolates, respectively. Its identification in a large number of isolates that test ETH-R indicates its role in resistance, in spite of it being found in a large number of isolates that test susceptible. A substitution in the neighboring position, N379D, was found in 13 and 40 solo R and solo S isolates, respectively.

The change from asparagine to aspartate is a change of charge, but retains the dimension of the amino acid, possibly leading to a small effect on ETH activation as compared to the change from a proline to a leucine in P378L. In other words, the nature of the mutations could influence the test result and contribute to ambiguity in their role in resistance. Indeed, in the 2023 WHO Catalogue P378L was removed from the Associated with Resistance category, while N379D raised to this category. Thus, the results here expand current understanding about the structural relevance of certain EthA residues as well as how the DSTs may be affected by modest MIC alterations caused by mutations.

## Ethics approval and consent to participate

Not applicable.

## Acknowledgements

We thank Rafael Ferreira for discussions and helpful suggestions. Sanger sequencing was carried out in the Fiocruz Genomics Platform (RPT01A).

