## Supplementary Figures for "In silico structural analysis of EthA substitutions for ranking priority mutations leading to ethionamide resistance in *Mycobacterium tuberculosis*"

**Figure S1.** Distribution of substitutions in the EthA amino acid sequence in 2831 *M. tuberculosis* clinical isolates as reported by Walker and colleagues (2022). Each dot represents a substitution found as (a and c) a solo R or S mutation in EthA or (b and d) in an isolate that has other mutations in EthA or other ETH resistance regions. Lower panels show the same data as upper panels, but for substitutions found in up to 40 isolates. Plotted with data from Table S1. The hotspots in Figure 1 are highlighted below each graph .

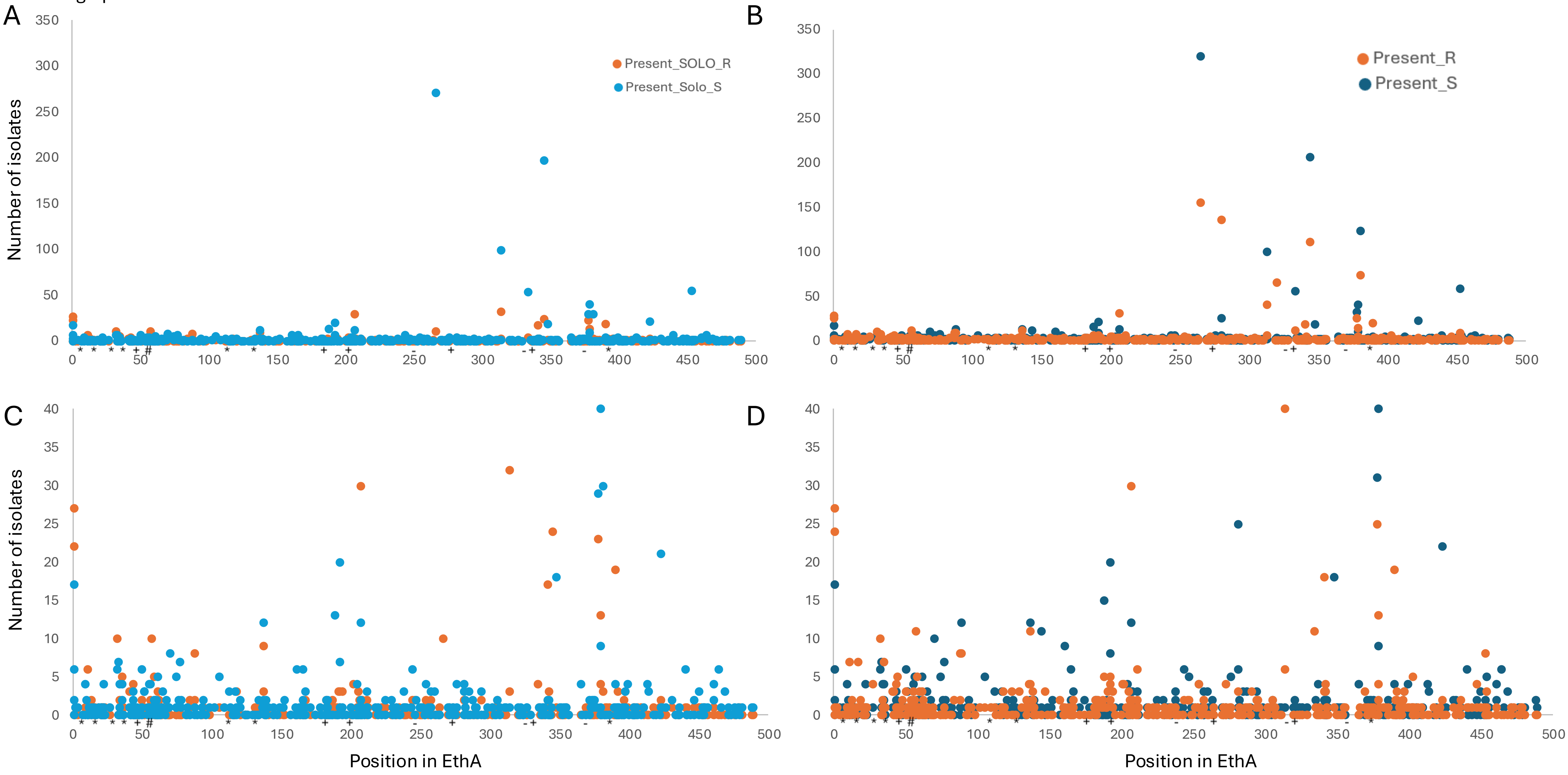

**Figures S2.** Distribution of EthA substitutions found as solo R mutations in at least 3 ETH-R *M. tuberculosis* clinical isolates, as reported by Walker and colleagues (2021). (a and c) Solo R mutations found in at least 3 isolates are shown as orange dots and when they are also found in Solo S isolates these are shown as light blue dots. (b and d) Isolates bearing these substitutions as well as other mutations in EthA or other ETH resistance genes. Lower panels show the same data as upper panels, but for substitutions found in up to 40 isolates. Plotted with data from Table S1. The hotspots in Figure 1 are highlighted below each graph .

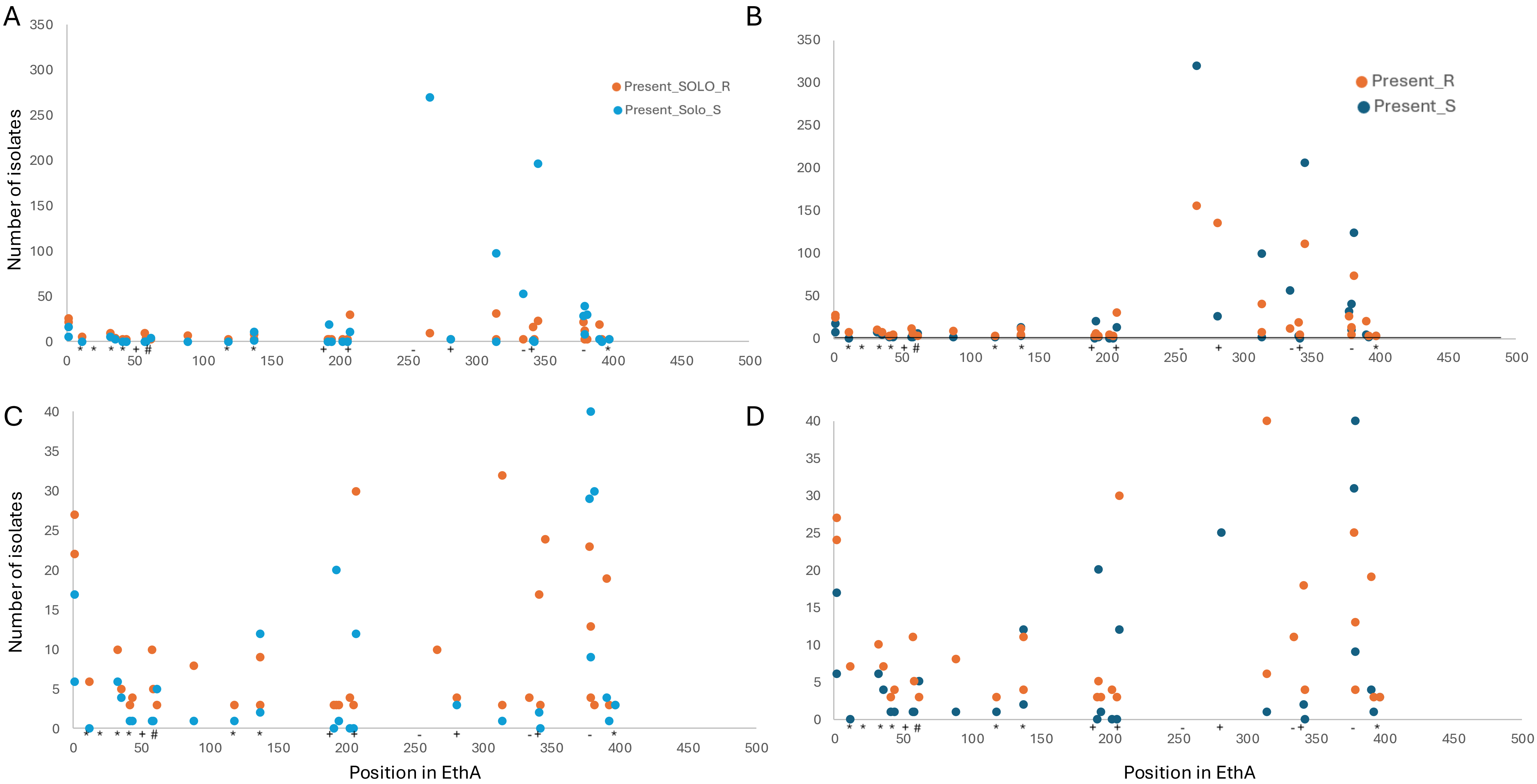

**Figure S3. 2D represenation of EthA - FAD interactions, generated with Maestro.**

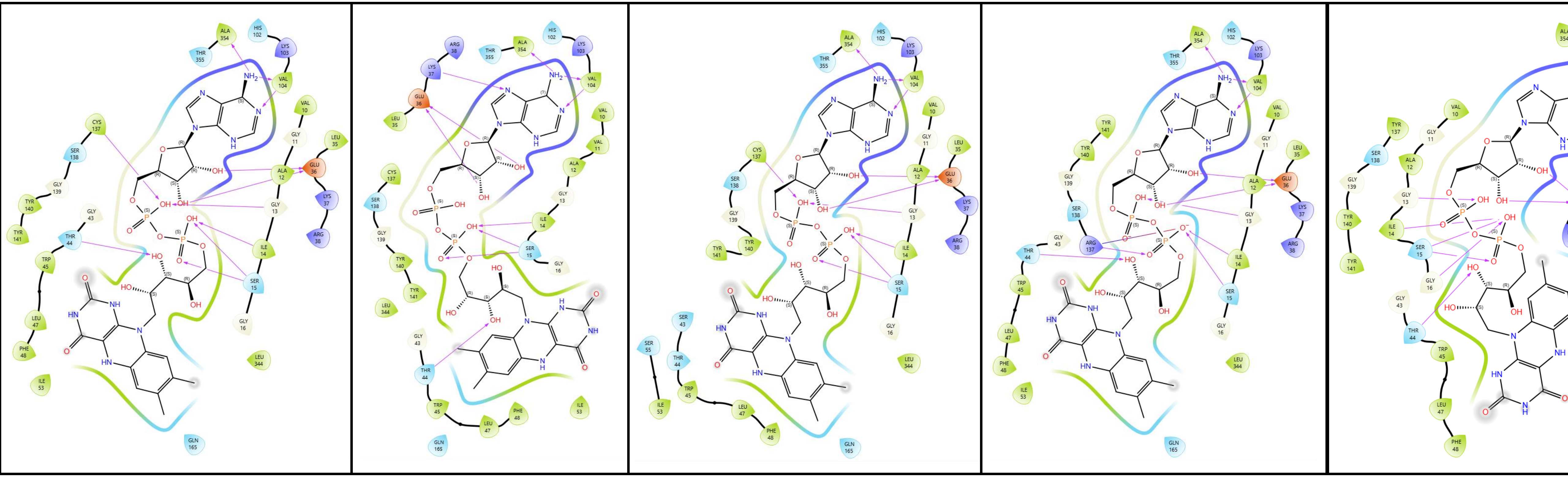

wt G11V G43S C137R C137Y

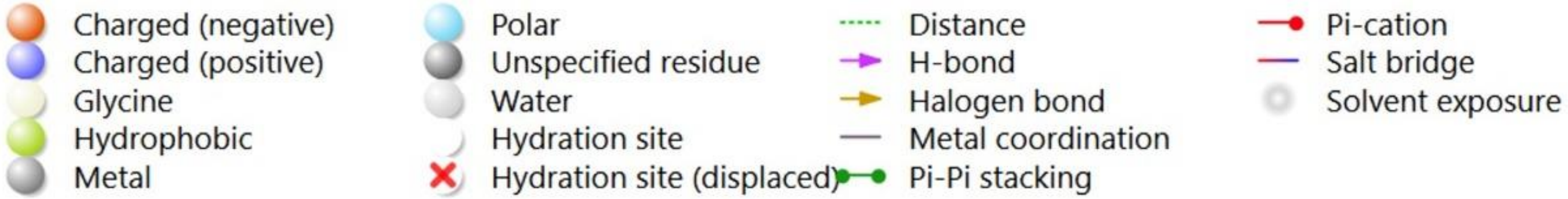

**Figure S4. 2D represenation of EthA - NAD interactions, generated with Maestro.**

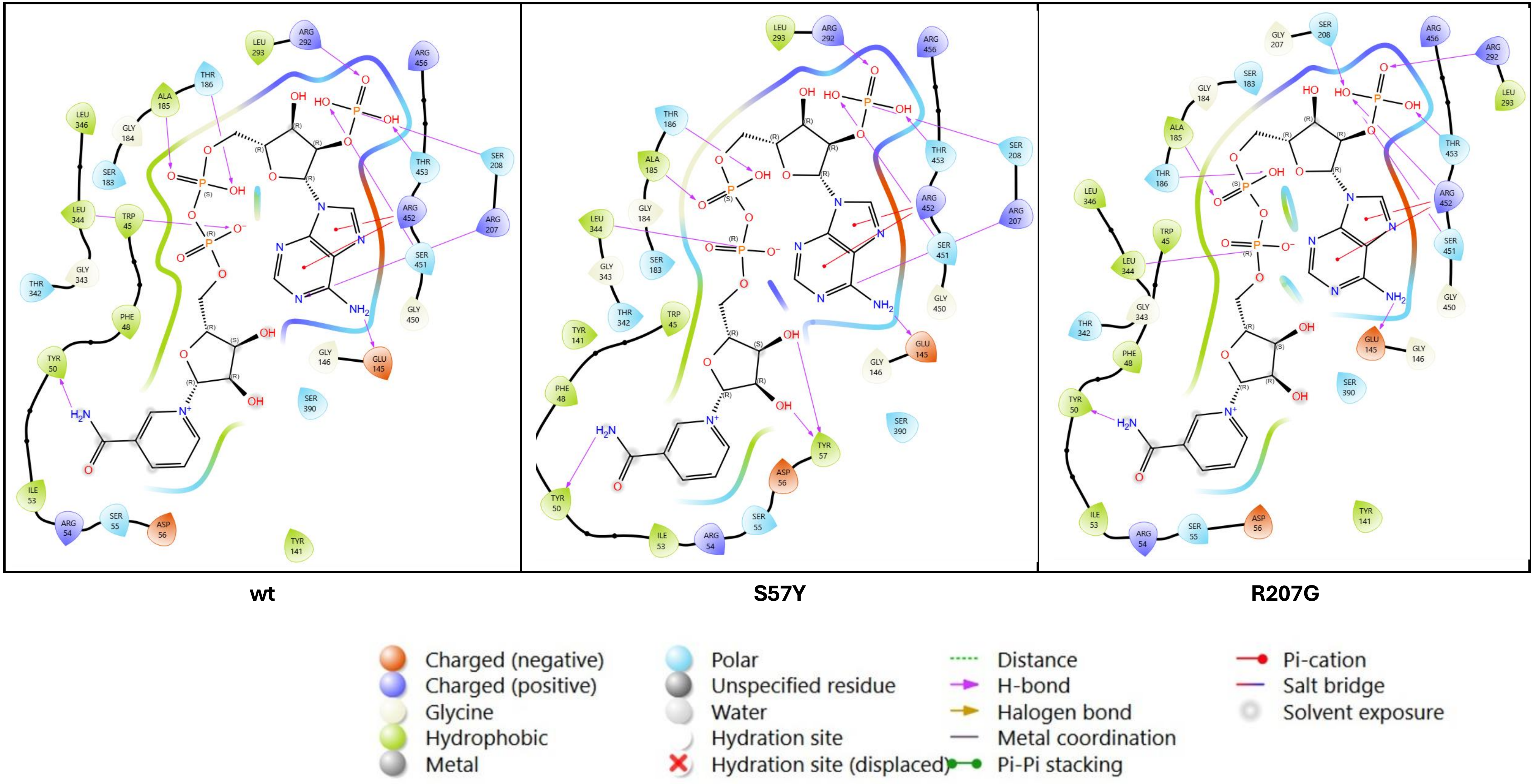

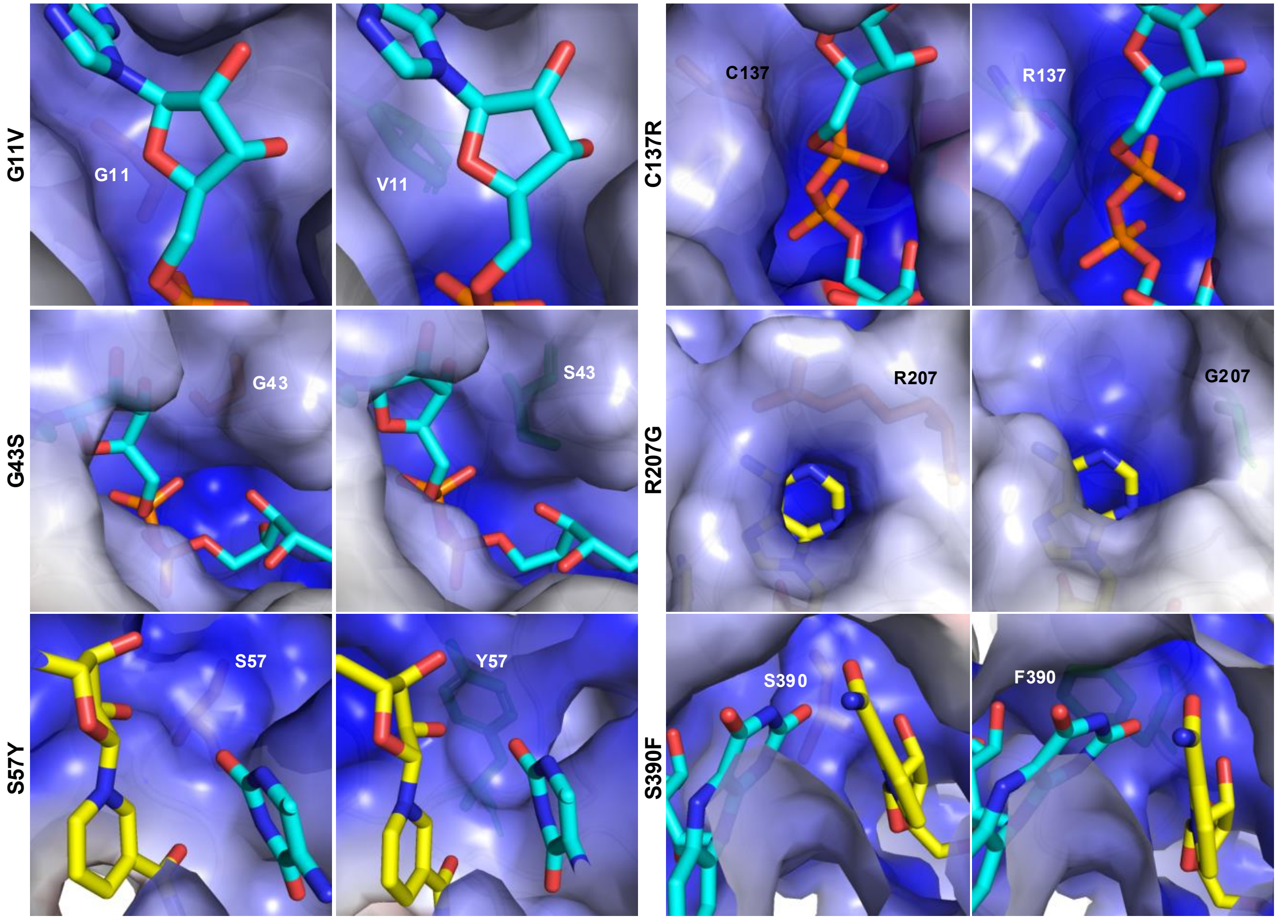


**Figure S5:** Mapping electrostatic potential surfaces on mutation sites G11V, G43S, S57Y, C137R, R207G, and S390F. The surface charge distributions were calculated using the APBS PyMOL plug-in. Charges vary between $-10^4$ (red) and $10^4$ (blue) eV.
